# Community detection in empirical kinase networks identifies new members of signaling pathways

**DOI:** 10.1101/2022.08.04.502751

**Authors:** Celia De Los Angeles Colomina Basanta, Marya Bazzi, Maruan Hijazi, Conrad Bessant, Pedro R. Cutillas

**Affiliations:** Cell signaling and Proteomics Group, Centre for Genomics and Computational Biology, Barts Cancer Institute, Queen Mary University of London, London EC1M 6BQ, UK; Warwick Mathematics Institute, University of Warwick, Coventry CV4 7AL, UK; The Alan Turing Institute, London NW1 2DB, UK; School of Biological and Chemical Sciences, Queen Mary University of London, London E1 4DQ, UK

## Abstract

Phosphoproteomics allows one to measure the activity of kinases that drive the fluxes of signal transduction pathways involved in biological processes such as immune function, senescence and growth. However, deriving knowledge of signaling network circuitry from these data is challenging due to a scarcity of phosphorylation sites that define kinase-kinase relationships. To address this issue, we previously identified around 6,000 phosphorylation sites markers of kinase-kinase relationships (that may be conceptualised as network edges), from which empirical cell-model-specific weighted kinase networks may be reconstructed. Here, we assess whether the application of community detection algorithms to such networks can identify new components linked to canonical signaling pathways.Phosphoproteomics data from acute myeloid leukaemia (AML) cells treated separately with PI3K, ATK, MEK and ERK inhibitors were used to reconstruct individual kinase networks. In each network, we applied the community detection method modularity maximization and selected the community containing the main target of the inhibitor the cells were treated with. These analyses returned communities that contained known canonical signaling components. Interestingly, in addition to canonical PI3K/AKT/MTOR members, the community assignments returned TTK (also known as MPS1) as a likely component of PI3K/AKT/mTOR signaling. We confirmed this observation with wet-lab laboratory experiments showing that TTK phosphorylation was decreased in AML cells treated with AKT and MTOR inhibitors. This study illustrates the application of community detection algorithms to the analysis of empirical kinase networks to uncover new members linked to canonical signaling pathways.

**Author summary:** Kinases are key enzymes that regulate the transduction of extracellular signals from cell surface receptors to changes in gene expression via a set of kinase-kinase interactions and signalling cascades. Inhibiting hyperactive kinases is a viable therapeutic strategy to treat different cancer types. Unfortunately, kinase signalling networks are robust to external perturbations, thus allowing tumour cells to orchestrate mechanisms that compensate for inhibition of specific kinases. Therefore, there is a need to better understand kinase network structure and to identify new therapeutic targets. Here, we reconstructed kinase networks from phosphoproteomics data, and compared the activity of its kinase interactions in acute myeloid leukaemia (AML) cells. We then tested community detection algorithms to identify kinase components associated to PI3K/AKT/MTOR signalling, a paradigmatic oncogenic signalling cascade. We found that TTK was usually grouped with networks derived for PI3K, AKT and MTOR kinases. Wet-lab experiments confirmed that TTK is likely to act downstream of AKT and MTOR. We thus show that our methods can be used to identify potential new members of canonical kinase signalling cascades.

## Introduction

Cells respond to changes in their environment through kinase-driven biochemical pathways that regulate the flux of signaling from cell surface receptors to changes in the expression levels of genes involved in cell functions [1]. Signal transduction is therefore essential to the upkeep of many biological processes including immune function [2], ageing/senescence [3], and growth, amongst others. Conversely, the dysregulation of signaling pathways can result in a range of different pathologies, such as metabolic syndrome, autoimmune diseases, or cancer [4,5]. Sustained proliferative signaling brought about by genetic mutations or by aberrant expression or activation of tumor suppressor genes and proto-oncogenes is a key cancer hallmark. For this reason, kinases are one of the most widely pursued targets for cancer therapeutics.

Although approaching signaling as a set of pathways is useful to conceptualize some of its properties, it is now recognized that signaling pathways form complex networks of interactions and enzymatic reactions [6]. A network in its simplest form is a graph, where nodes represent entities of interest and edges represent pairwise interactions between such nodes [7]. Quantitative phosphoproteomics can provide new insights into these complex kinase signaling networks through the measurement of kinase phosphorylation, and capture the complexity of interactions within and between kinase signaling pathways at the network level [8,9]. While quantitative phosphoproteomics have been widely used to study changes in kinase activities in cancer cells [10–14], studies that incorporate quantitative phosphoproteomics with kinase-substrate networks are fewer and relatively recent [8,9,15–17]. A problem has been the lack of phosphorylation sites that are markers of kinase network circuitry and which therefore allow for cell-type specific reconstruction of signaling networks from phosphoproteomics data. To address this issue, Hijazi *et al* recently described a set of 6,000 phosphorylation sites that define kinase-kinase interactions [16], from which networks may be reconstructed in a given cellular state.

Here, we aimed to characterize novel kinase interactions in canonical cancer signaling pathways. To do so, we construct networks of kinase-kinase relationships with edge weights defined by the change in the levels of markers of kinase-kinase relationships in response to the inhibition of a given kinase (as measured by quantitative phosphoproteomics). Once networks are constructed for each perturbation, we applied the community detection approach modularity maximization, which has been used in a wide variety of biological networks [18] and has achieved favorable performance in comparative studies on biological networks [19]. In doing so, our aim is to illustrate how one can use community detection techniques from network science to analyse signaling derived from quantitative phosphoproteomics, and to demonstrate that studying the mesoscale structure (e.g., community structure) [7,20] of the resulting network can reveal novel biological insights about cancer kinase signaling pathways.

## Materials and methods

### Data description

We first analyzed enrichment values of 1500 kinase-kinase interactions in P31/FUJ cells treated separately with the kinase inhibitors trametinib, GDC0994, GDC0941 and AZD5363, measured by z-score analysis as in the Kinase Substrate Enrichment Analysis (KSEA) method [10] (these values were taken from four of the columns in the Supplementary dataset 5 published in [16]). The kinase-kinase interactions were defined in previous work [16] based on the existence of common putative downstream targets (PDTs), i.e. both kinases in a kinase-kinase relationship that define a given edge act upstream of a number of phosphorylation sites. We focus on kinase-kinase relationships inhibited by the treatments (i.e., with negative z-score values) where one has some notion of ground truth for the pathway containing the main target of the inhibitor. That is, MAP2K1 and MAPK1/3, the respective targets of trametinib and GDC0994, belong to the MEK/ERK pathway [21], while PIK3CA and AKT1/2, the respective targets of GDC0941, AZD5363, belong to the PI3K/AKT/mTOR pathway [3]. Table 1 shows summary statistics for the dataset portion that we focus on in the present paper. This paper focuses on the analysis of the GDC0941 and AZD5363 treatments, while that of trametinib and GDC0994 treated cell measurements can be found in the Supplementary Information.

**Table 1.** Summary statistics for strictly negative z-scores.

| Treatment | No. of kinases | min | max | $\mu$ | $\sigma$ |
| --- | --- | --- | --- | --- | --- |
| trametinib | 69 | -3.732 | -0.000607 | -0.278 | 0.276 |
| GDC0941 | 72 | -6.749 | -0.00178 | -0.590 | 0.779 |
| AZD5363 | 74 | -16.886 | -0.00334 | -0.505 | 0.966 |
| GDC0994 | 67 | -1.411 | -0.00233 | -0.309 | 0.261 |

Data available in [16] (see Data description). The column “min” (resp., “max”) corresponds to the smallest (resp., largest) strictly negative z-score for the corresponding treatment. The columns $\mu$ and $\sigma$ correspond to the mean and standard deviation, respectively, of strictly negative z-score values for each treatment

### Network construction

In this study, we work under the hypothesis that the inhibition of a kinase that belongs to a pathway will result in the inhibition of the kinase interactions within that pathway. Therefore we only consider the kinase interactions with negative z-scores (i.e., inhibited) for each treatment, from which we construct four weighted and undirected networks (i.e., without assumption on whether the kinases act upstream or downstream of each other). We adopt this approach because the presence of a given edge (kinase-kinase interaction) in the network would be revealed by the decrease in phosphorylation sites that define such edges as a result of treatment with specific kinase inhibitors.

Loosely speaking, a community in a network is a set of nodes that are “more densely” connected to each other than they are to nodes in the rest of the network [20]. Since we are interested in capturing the kinase interactions with the highest decrease in activity (i.e., lowest z-scores) in response to a treatment, we use the absolute value of z-scores as the weights of the kinase-kinase edges. The weights of the kinase-kinase edges is then proportional to the extent of inhibition that the kinase-kinase interaction it represents experiences with said treatment (compared to control).

### Community detection

We denote by *n* the number of nodes in a network and by ***A*** the corresponding adjacency matrix. In this paper, the nodes are kinases and the entries of ***A*** are the absolute value of negative z-scores (see Network construction). We use *modularity maximization* to identify communities. The modularity maximization problem can be stated as follows:

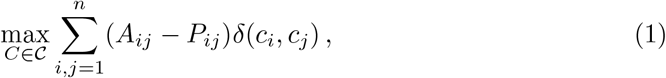

where 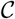 is the set of all n-node partitions, ***A*** is the adjacency matrix of the observed network, ***P*** is an expected null network under some null model, and *c_i_* is the set assignment of node *i* [20,22]. This method partitions a network into sets of nodes called “communities” that have a larger total internal edge weight than that expected in the same sets in a null network, generated from some null model.

To find locally optimal partitions for the community detection approach modularity maximization in Community detection, we use the locally greedy algorithmic heuristic Louvain [23]. We consider both a “uniform” and “Newman-Girvan” null network [24], and choose the uniform null network for our experiments on the basis of slightly better alignment with “ground truth” canonical pathways (see Results) across different algorithmic runs. We use a nondeterministic version of Louvain in which node order is randomized at the start of each iteration. This stochasticity can yield different partitions across different runs. To compute a “consensus partition”, we use the iterative approach in [25]. We obtain an ensemble of partitions using the Louvain algorithm (the size of the ensemble is the number of algorithmic runs), for which we compute a “co-classification matrix”. We reiterate our community detection procedure on the co-classification matrix until the new co-classification matrix resulting from the corresponding partition ensemble is binary (typically after 1 or 2 iterations). The partitions we obtain are robust (e.g., to different repetitions of the consensus procedure), see S1 Appendix. We show adjacency matrices and corresponding co-classification matrices in fig. 1.

**Fig 1.**
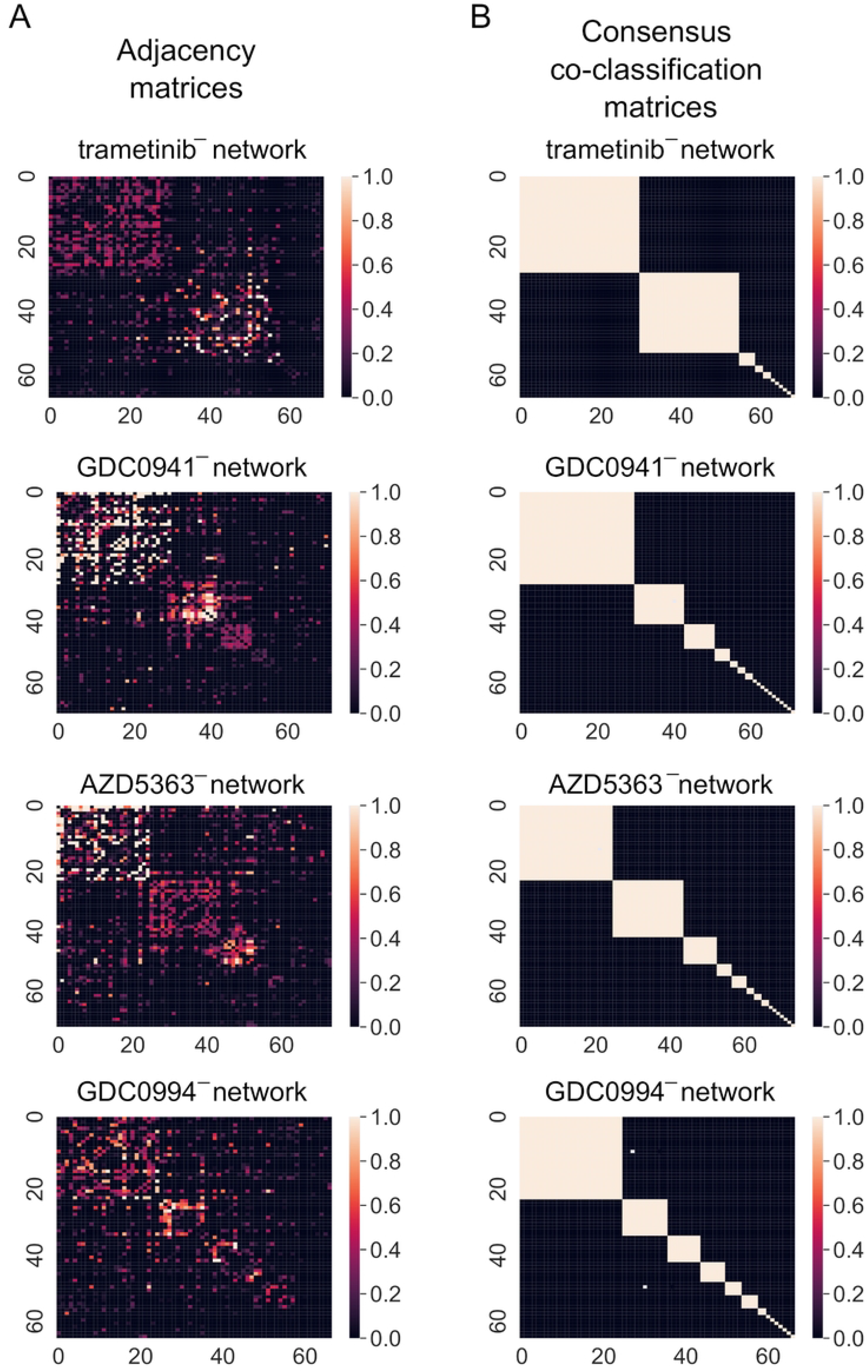
Results of community detection from phosphoproteomics-derived networks. Adjacency matrix (panel A) and consensus co-classification matrix (panel B). In panel A, we show the adjacency matrix of each kinase network ordered by their community assignment in the consensus partition. We cap the entries of the adjacency matrix at 1 to emphasize structure. In panel B, we show the co-classification matrix of a consensus partition (see Community detection), which takes the values 1 or 0. The colors scale with the entries of the matrices.

### Wet-lab experiments: phosphoproteomics analysis of kinase inhibitor-treated cells

To experimentally verify novel computational findings, phosphoproteomics analysis was carried out as described in previous work [16,26]. Briefly, the P31/FUJ cell line was grown in RPMI medium supplemented with 10% FBS and 1% Pen/Strep. Cells were treated with 1 μM CFI-402257 (TTKi), MK2206 (AKTi), BYL719 (PI3Ki) or AZD8055 (MTORi) for three hours. Subsequently, cells were lysed in urea buffer and digested with trypsin. Finally, phosphopeptides were enriched using TiO2 and analysed in a LC-MS/MS system, which consisted of an Ultimate 3000 ultra-high-pressure chromatography connected to a Q-Exactive Plus mass spectrometer. Peptide identification and quantification was performed using Mascot search engine and Pescal, respectively, as described in [26]. The mass spectrometry proteomics data has been deposited to the ProteomeXchange Consortium via the PRIDE partner repository [27] with the dataset identifier PXD026039.

## Results

### Community structure can reflect kinase signaling pathways

The main aim of this study is to assess whether the application of community detection techniques to quantitative phosphoproteomics based kinase networks can identify new members of canonical kinase signaling pathways. For this purpose, we first assess whether known kinases in the signaling pathway PI3K/AKT/mTOR belong to the same community. That is, we compare the results of community detection with what we regard as “ground truth” in the present context.

Two networks were constructed based on the edge enrichment of kinase-kinase relationships (see Network construction), measured as z-scores, of AML cells treated separately with the kinase inhibitors GDC0941 (a PIK3CA/PI3K inhibitor) and AZD5363 (AKT inhbitor). We refer to these networks as GDC0941^-^ and AZD5363^-^, respectively, for the remainder of the paper, where the superscript “^-^” is to emphasize that we only consider negative z-scores (i.e., kinase-kinase relationships that were inhibited by the compounds). Furthermore, in each network we only focus on the community containing the main target of the inhibitor the cells were treated with. For example, the communities containing PI3K kinase alpha isoform (gene name PIK3CA) and AKT1/2^1^ were selected for networks GDC0941^-^ and AZD5363^-^, respectively, and we denote these respective communities by 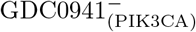 and 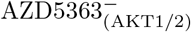.

PIK3CA and AKT1/2 are two of the main members of the PI3K/AKT/mTOR signaling pathway, which is often activated in a range of cancers and contributes to their development [28]. Thus, we expect the selected communities in networks GDC0941^-^ and AZD5363^-^ to be similar or identical in terms of content, and a reflection of the kinase interactions in this pathway. We found that the communities GDC0941 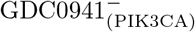 and 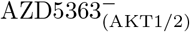 share 24 kinases (blue nodes in fig. 2A and fig. 2B) while CDK2 is part of the community 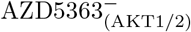 (purple node in fig. 2A) but not 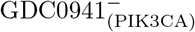. Six other kinases were part of 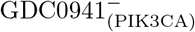 but not 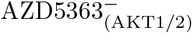 (purple nodes in fig. 2B), four of which we attribute to MAP2K1 signaling (further discussed in S2 Appendix). Of note, PIK3CA, AKT1/2,mTOR and RPS6KB1, which are well known members of the canonical PI3K/AKT/mTOR signaling pathway [28], are present in both communities, which is concomitant with the fact that the edges between these kinases have large weight values (see heatmaps in panel A and B of Figure 2) in both networks (i.e. their activity is highly decreased in response to both inhibitors).

**Fig 2.**
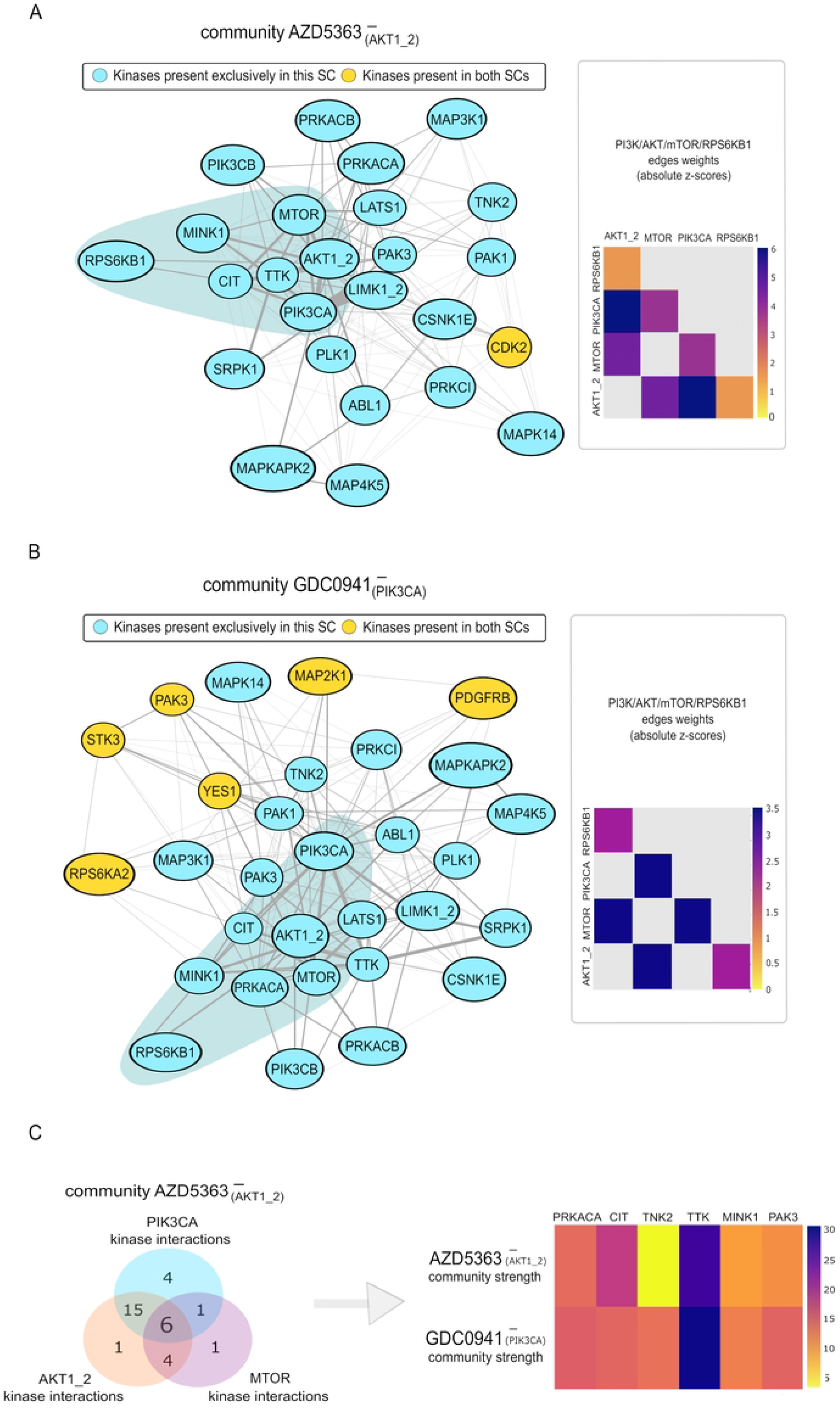
Identification of Canonical and non-canonical kinases in PI3K/AKT networks. A.Kinase interactions of the selected communities in network AZD5363^-^. Abbreviations: SC; selected community. On the left hand side of panel A the nodes and edges within the community 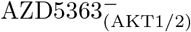 are depicted, in which the yellow nodes represent kinases present in 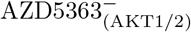 but not 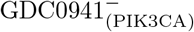, and the blue ones represent the nodes present in both communities. B. Kinase interactions of the selected communities in network GDC0941^-^. On the left hand side of panel B the nodes and edges within the community 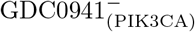 are depicted, in which the yellow nodes represent kinases present in 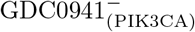 but not in 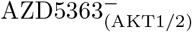, and the blue ones represent the nodes that are present in both communities. The heatmaps on the right hand of panel A and B show the weights of the edges between well known members of the PI3K/AKT/mTOR pathway in each of the communities, in which a grey cell indicates that interaction is not present in that community. C.Kinase interactions and community strength show TTK is likely to be downstream of PI3K/AKT/mTOR signaling. Panel A shows the intersections between the PIK3CA, AKT1/2 and MTOR edges/kinase interactions in the 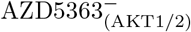 community, in which six kinases have direct interactions with PIK3CA, AKT1/2 and MTOR. In panel B, the community strength (i.e. the sum of the weight of the edges between a node and other nodes in its community) of each of the said six kinases in the 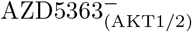 and 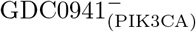 communities is depicted.

Thus, the community detection algorithm used in this study returned the PI3K/AKT/MTOR canonical signaling pathway, suggesting that this approach can identify biologically meaningful associations. We stress, however, that community detection alone is hypothesis generating and not sufficient for the unambiguous identification of novel members in said pathways, for which we delineate further steps in the next two sections.

### TTK is a likely downstream target of PI3K/AKT/mTOR signaling

We next investigated which of the kinases placed in the communities 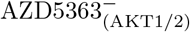 and 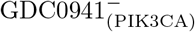 are most likely to act downstream of PI3K/AKT/mTOR signaling. To do so, we narrowed the list of 24 kinases assigned to both the 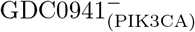 and 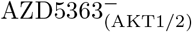 community (see Community structure can reflect kinase signaling pathways) based on two criteria. First, we discarded all kinases that do not have a direct edge with all three kinases PIK3CA, AKT1/2 and mTOR in the AZD5363^-^ network. If a kinase does not have a direct relationship with all three kinases in PI3K/AKT/mTOR signaling (see Network construction) it is more likely to act downstream of some but not all kinases, since it either means that the kinase does not share downtream targets with PI3K, AKT and mTOR, or that the relationship is not included in the network because its activity is unchanged or increased in response to the different inhibitors. We found that, in addition to the known PI3K/AKT/MTOR pathway members AKT1/2, mTOR and RPS6KB1, six kinases have direct edges with PIK3CA, AKT1/2 and mTOR in the 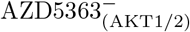 community, as shown in fig. 2C. Secondly, we computed the *community strength* (CS) (i.e., the sum of the weight of the edges between a node and other nodes in its community) of said kinases in 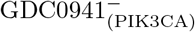 and 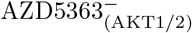 (see fig. 2C). Interestingly, TTK had the highest CS by far out of the six selected kinases in both communities, as well as the third highest CS of all kinases in 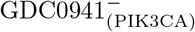 and the fourth highest CS of the 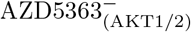 kinases, meaning its interactions with the other kinases in both communities have the highest decrease in activity in response to both PIK3CA and AKT1/2 inhibition. These results strongly suggest that TTK is likely associated to PI3K/AKT/mTOR signaling.

### Phosphoproteomics analysis confirms that TTK is downstream of PI3K/AKT/mTOR signaling

As shown in the previous section, the kinase TTK – a mitotic kinase also known as MPS1 – was found to be in the same community as PI3K, AKT and MTOR, suggesting that TTK may be upstream or downstream of the PI3K/AKT/MTOR pathway. To validate this finding, the cell line P31/FUJ was treated with 1 μM CFI-402257 (a highly specific small molecule inhibitor of TTK [29], thereafter named TTKi), MK2206 (AKTi), BYL719 (PI3Ki) or AZD8055 (MTORi) for 3h, and further subjected to phosphoproteomics analysis by mass spectrometry, as in previous work [16]. This analysis led to the identification and quantification of more than 8,000 phosphopeptides in biological and technical replicate (fig. 4A). As volcano plots in Figure 1B show, at the same threshold of statistical significance (unadjusted *p*-value < 0.01), the MTORi caused the highest impact on the phosphoproteome (577 increased and 525 decreased phosphopeptides), whereas the TTKi only increased 59 and decreased 89 phosphosites compared to the control condition (fig. 3B). Treatments with PI3Ki and AKTi impacted more than 400 phosphorylation sites each. The larger impact on protein phosphorylation of PI3K/AKT/MTOR compared to TTK inhibition, suggests that TTK is unlikely to be upstream of the PI3K pathway.

**Fig 3.**
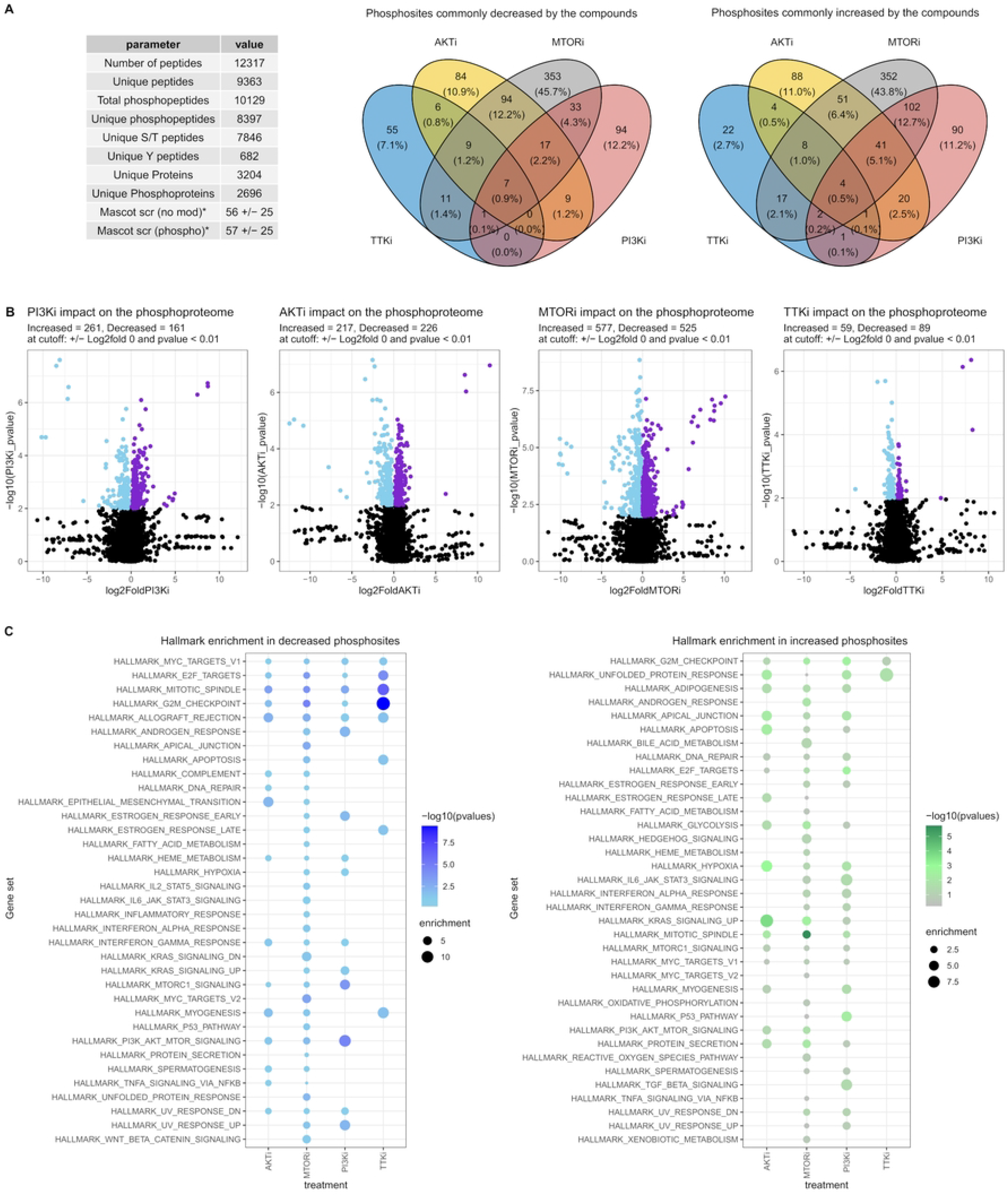
TTK and PI3K/ATK/mTOR pathway inhibitors impact common phosphorylation pathways. A. Summary of results of phosphoproteomics experiment. Venn diagrams show phosphopeptides commonly decreased or increased by the kinase inhibitors. B. Impact on the phosphoproteome as a function of the kinase inhibitors. Phosphopeptides increased or decreased significantly are highlighted in purple or sky blue, respectively. P-values were calculated by t-test of log transformed data. C. Enrichment of hallmark gene sets from the Molecular Signature Database (MSigDB) considering phosphopeptides decreased (left panel) or increased (right panel) as a function of the compound treatments.

**Fig 4.**
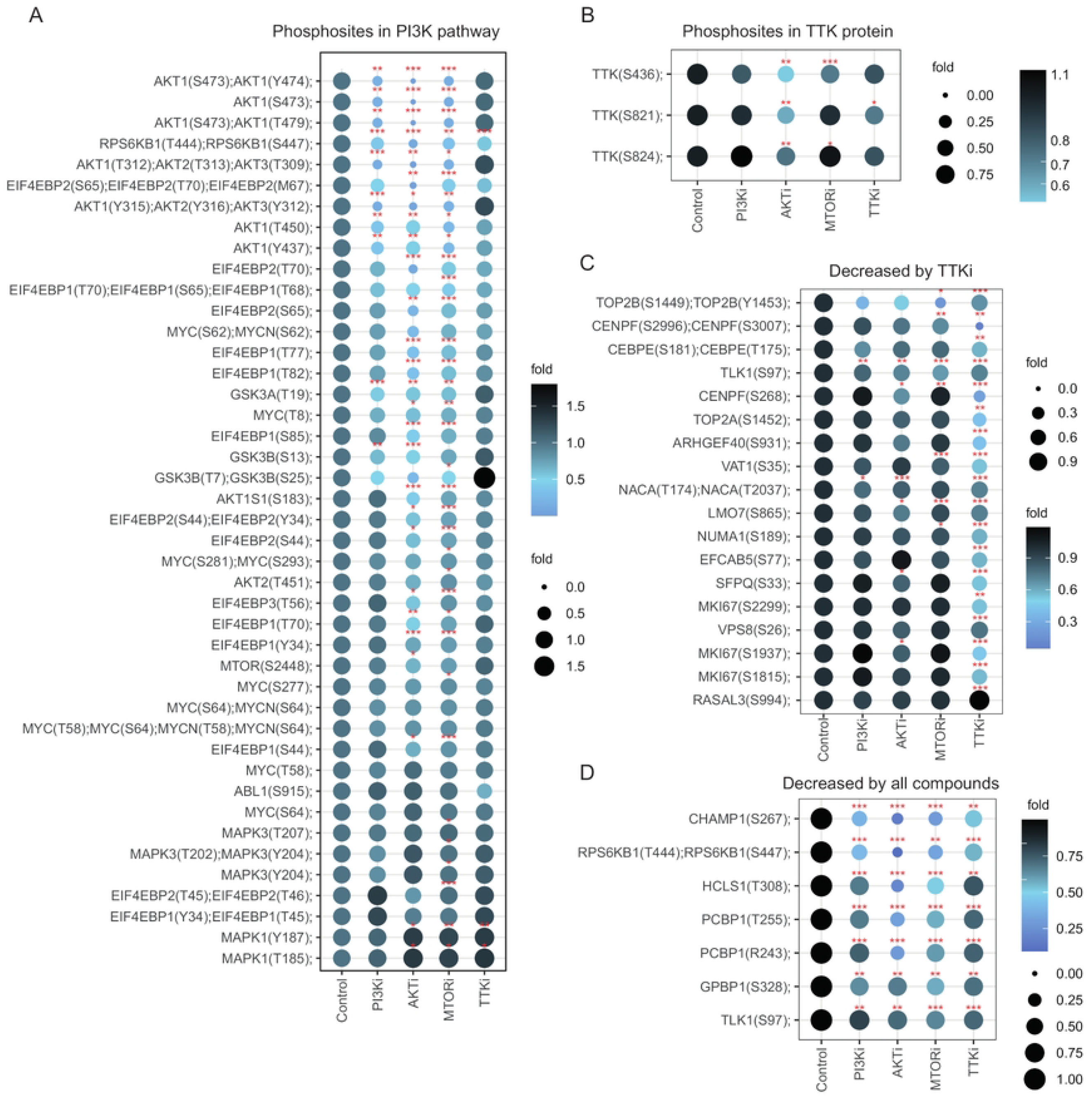
Phosphoproteomics of cells treated with kinase inhibitor supports a link between TTK and PI3K/AKT/mTOR signalling. Dot plots denote relative intensities of phosphosites as a function of treatment with the named kinase inhibitors, grouped by those linked to PI3K pathway activity (A), on TTK (B), decreased exclusively by the TTK inhibitor (C) or commonly decreased across all the treatments (D). p-values were calculated by t-test. \**p* < 0.05, * * *p* < 0.01, * * \**p* < 0.001, n = 4.

We found 7 and 4 phosphorylation sites commonly decreased or increased by all the kinase inhibitors, respectively (fig. 4A). Across the 7 phosphosites decreased, we highlight the double phosphorylated RPS6KB1 at T444 and S447, located in an autoinhibitory domain [30], the CDK1-mediated phosphorylation of CHAMP1 required for the attachment of spindle microtubules to the kinetochore [31] and the inhibition by phosphorylation of TLK1, which follows the generation of DNA double-stranded breaks during S phase as a DNA damage checkpoint (fig. 4D) [32]. These observations are consistent with the known roles of the PI3K pathway and the TTK mitotic kinase in the cell-cycle regulation and indicate a functional relationship between the two.

Furthermore, several markers of PI3K pathway activity, including multiple phosphorylation sites in AKT1/2, 4EPB1/2, PRAS40, GSK3A/B, MYC, were inhibited by PI3Ki, AKTi, MTORi but not by TTKi (fig. 4A), implying that TTK is not directly upstream of the PI3K pathway. Interestingly, we also found that the AKTi and MTORi significantly decreased the phosphorylation of TTK at S824 and S436 (fig. 4B), whose function are to regulate the activity of TTK in controlling cytoskeletal reorganization [33]. The TTKi only affected the phosphorylation at S821, which implies changes in kinetochore localization [34]. Moreover, a pathway analysis using hallmark gene sets revealed an enrichment of the PI3K/AKT/MTOR signaling in phosphoproteins that decreased as a result of PI3Ki, AKTi or MTORi treatment, but not by TTKi (fig. 4C). In contrast, hallmark genes with roles in G2M checkpoint and mitotic spindle were enriched in phosphoproteins decreased by TTK as well as PI3K pathway inhibitors. Taken together, these observations are consistent with the finding obtained with the community detection algorithm and suggest that TTK phosphorylation and at least some TTK functions are downstream of the PI3K/AKT/MTOR network.

## Discussion

Studying the effects of kinase pathways dysregulation and over-activation on carcinogenesis has led to a better understanding of cancer biology and the development of therapies that target oncogenic kinase signaling. However, intrinsic and acquired resistance to treatments with kinase inhibitors and other drugs, which occur in most patients, is estimated to be responsible for 90% of cancer deaths [35]. Acquired resistance may occur through target modification, in which the drug target acquires further mutations that result in reduced drug binding, thus reducing the effectiveness of the treatment [36] or by the rewiring of kinase networks so that cancer cells use parallel or downstream pathway to proliferate [37]. The discovery of new members of oncogenic pathways, such as that of TTK in the PI3K/AKT/mTOR pathway in this study, may help the identification of alternative targets for cancer treatment.

The two main aims of this study were first to assess whether it is possible to define sub-clusters of kinases that represent individual kinase signaling pathways by applying community detection algorithms to networks derived from empirical markers of kinase-kinase activity, and secondly, to assess whether such an approach can be used to identify new kinase members associated to canonical signaling pathways. Our study differs from most previous works on quantitative phosphoproteomics-based kinase network analysis, which focused on the prediction of kinase interactions using publicly available kinase-substrate databases, such as PhosphoSitePlus, Signor or Phospho.ELM (which draw information from systematic literature mining) [10,14]. A prior study [15] focused on identifying kinases that mediate crosstalk between different signaling pathways downstream of the activation of a receptor kinase. To do so, Narushima *et al* investigated a kinase-substrate network where interactions between distinct kinases are unweighted. Our study shows that, by harnessing newly available large scale phosphoproteomics datasets [16], community detection algorithms may also be used to identify new members of canonical signaling pathways.

Overall, our study demonstrates that the study of community structure has the potential to reveal novel biological insights about cancer kinase signaling pathways. We found that (1) community assignments align well with established canonical pathways and (2) communities can reveal new components of known pathways that may be validated in wetlab experiments.

Future research in this field could include the application of network science techniques to investigate further questions in cell signalling, such as the relationship between kinase mesoscale network structure and an individual’s response to cancer treatment. Ultimately, the hope is that such research can improve our understanding of biochemical pathways and, as a result, improve methods for drug response prediction.

## Supporting information

**S1 Appendix. Partition and community robustness.**

**S2 Appendix. Analysis of kinase communities with strong links to MAP2K1 and MAPK1/3.**

## Footnotes

1 AKT isoforms 1 and 2 are considered as one since they have the same z-scores due to having the same putative downstream targets assignments.

## References

1. Schlessinger J. Cell Signaling by Receptor Tyrosine Kinases. Cell. 2000;103(2):211–225. doi:10.1016/s0092-8674(00)00114-8.

2. Ross SH, Cantrell DA. Signaling and Function of Interleukin-2 in T Lymphocytes. Annual Review of Immunology. 2018;36(1):411–433. doi:10.1146/annurev-immunol-042617-053352.

3. Tan P, Wang YJ, Li S, Wang Y, He JY, Chen YY, et al. The PI3K/Akt/mTOR pathway regulates the replicative senescence of human VSMCs. Molecular and Cellular Biochemistry. 2016;422(1-2):1–10. doi:10.1007/s11010-016-2796-9.

4. Banerjee S, Biehl A, Gadina M, Hasni S, Schwartz DM. JAK–STAT Signaling as a Target for Inflammatory and Autoimmune Diseases: Current and Future Prospects. Drugs. 2017;77(5):521–546. doi:10.1007/s40265-017-0701-9.

5. Hanahan D, Weinberg RA. Hallmarks of Cancer: The Next Generation. Cell. 2011;144(5):646–674. doi:10.1016/j.cell.2011.02.013.

6. C Jørgensen RL. Simplistic pathways or complex networks? Current Opinion in Genetics & Development. 2010;20(1):15–22. doi:10.1016/j.gde.2009.12.003.

7. Newman MEJ. Networks. Oxford University Press; 2018.

8. Wilkes EH, Terfve C, Gribben JG, Saez-Rodriguez J, Cutillas PR. Empirical inference of circuitry and plasticity in a kinase signaling network. Proceedings of the National Academy of Sciences. 2015;112(25):7719–7724. doi:10.1073/pnas.1423344112.

9. Terfve CDA, Wilkes EH, Casado P, Cutillas PR, Saez-Rodriguez J. Large-scale models of signal propagation in human cells derived from discovery phosphoproteomic data. Nature Communications. 2015;6(1). doi:10.1038/ncomms9033.

10. Casado P, Rodriguez-Prados JC, Cosulich SC, Guichard S, Vanhaesebroeck B, Joel S, et al. Kinase-Substrate Enrichment Analysis Provides Insights into the Heterogeneity of Signaling Pathway Activation in Leukemia Cells. Science Signaling. 2013;6(268):rs6–rs6. doi:10.1126/scisignal.2003573.

11. Wandinger SK, Lahortiga I, Jacobs K, Klammer M, Jordan N, Elschenbroich S, et al. Quantitative Phosphoproteomics Analysis of ERBB3/ERBB4 Signaling. PLOS ONE. 2016;11(1):e0146100. doi:10.1371/journal.pone.0146100.

12. Robitaille AM, Christen S, Shimobayashi M, Cornu M, Fava LL, Moes S, et al. Quantitative Phosphoproteomics Reveal mTORC1 Activates de Novo Pyrimidine Synthesis. Science. 2013;339(6125):1320–1323. doi:10.1126/science.1228771.

13. T E Bass DC. Quantitative phosphoproteomics reveals mitotic function of the ATR activator ETAA1. Journal of Cell Biology. 2019;218(4):1235–1249. doi:10.1083/jcb.201810058.

14. Ochoa D, Jonikas M, Lawrence RT, El Debs B, Selkrig J, Typas A, et al. An atlas of human kinase regulation. Mol Syst Biol. 2016;12(12):888. doi:10.15252/msb.20167295.

15. Narushima Y, Kozuka-Hata H, Tsumoto K, Inoue JI, Oyama M. Quantitative phosphoproteomics-based molecular network description for high-resolution kinase-substrate interactome analysis. Bioinformatics. 2016;32(14):2083–2088. doi:10.1093/bioinformatics/btw164.

16. Hijazi M, Smith R, Rajeeve V, Bessant C, Cutillas PR. Reconstructing kinase network topologies from phosphoproteomics data reveals cancer-associated rewiring. Nature Biotechnology. 2020;38(4):493–502. doi:10.1038/s41587-019-0391-9.

17. Buljan M, Ciuffa R, van Drogen A, Vichalkovski A, Mehnert M, Rosenberger G, et al. Kinase Interaction Network Expands Functional and Disease Roles of Human Kinases. Molecular Cell. 2020;79(3):504–520.e9. doi:10.1016/j.molcel.2020.07.001.

18. Liu C, Ma Y, Zhao J, Nussinov R, Zhang YC, Cheng F, et al. Computational network biology: Data, models, and applications. Physics Reports. 2020;846:1–66. doi:10.1016/j.physrep.2019.12.004.

19. Choobdar S, Ahsen ME, Crawford J, Tomasoni M, Fang T, Lamparter D, et al. Assessment of network module identification across complex diseases. Nature Methods. 2019;16(9):843–852. doi:10.1038/s41592-019-0509-5.

20. Fortunato S, Hric D. Community detection in networks: A user guide. Physics Reports. 2016;659:1–44. doi:10.1016/j.physrep.2016.09.002.

21. McCubrey JA, Steelman LS, Chappell WH, Abrams SL, Wong EWT, Chang F, et al. Roles of the Raf/MEK/ERK pathway in cell growth, malignant transformation and drug resistance. Biochimica et Biophysica Acta (BBA) - Molecular Cell Research. 2007;1773(8):1263–1284. doi:10.1016/j.bbamcr.2006.10.001.

22. Newman MEJ, Girvan M. Finding and evaluating community structure in networks. Physical Review E. 2004;69(2). doi:10.1103/physreve.69.026113.

23. Blondel VD, Guillaume JL, Lambiotte R, Lefebvre E. Fast unfolding of communities in large networks. Journal of Statistical Mechanics: Theory and Experiment. 2008;2008(10):P10008. doi:10.1088/1742-5468/2008/10/p10008.

24. Bazzi M, Porter MA, Williams S, McDonald M, Fenn DJ, Howison SD. Community Detection in Temporal Multilayer Networks, with an Application to Correlation Networks. Multiscale Modeling & Simulation. 2016;14(1):1–41. doi:10.1137/15m1009615.

25. Lancichinetti A, Fortunato S. Consensus clustering in complex networks. Scientific Reports. 2012;2(1). doi:10.1038/srep00336.

26. Gerdes H, Casado P, Dokal A, Hijazi M, Akhtar N, Osuntola R, et al. Drug ranking using machine learning systematically predicts the efficacy of anti-cancer drugs. Nature Communications. 2021;12(1). doi:10.1038/s41467-021-22170-8.

27. Perez-Riverol Y, Csordas A, Bai J, Bernal-Llinares M, Hewapathirana S, Kundu DJ, et al. The PRIDE database and related tools and resources in 2019: improving support for quantification data. Nucleic Acids Research. 2018;47(D1):D442–D450. doi:10.1093/nar/gky1106.

28. Aoki M, Fujishita T. Oncogenic Roles of the PI3K/AKT/mTOR Axis. In: Current Topics in Microbiology and Immunology. Springer International Publishing; 2017. p. 153–189.

29. Mason JM, Wei X, Fletcher GC, Kiarash R, Brokx R, Hodgson R, et al. Functional characterization of CFI-402257, a potent and selective Mps1/TTK kinase inhibitor, for the treatment of cancer. Proceedings of the National Academy of Sciences. 2017;114(12):3127–3132. doi:10.1073/pnas.1700234114.

30. Weng QP, Kozlowski M, Belham C, Zhang A, Comb MJ, Avruch J. Regulation of the p70 S6 Kinase by Phosphorylation in Vivo. Journal of Biological Chemistry. 1998;273(26):16621–16629. doi:10.1074/jbc.273.26.16621.

31. Itoh G, Kanno S, Uchida KSK, Chiba S, Sugino S, Watanabe K, et al. CAMP (C13orf8, ZNF828) is a novel regulator of kinetochore-microtubule attachment. The EMBO Journal. 2011;30(1):130–144. doi:10.1038/emboj.2010.276.

32. Groth A. Human Tousled like kinases are targeted by an ATM-and Chk1-dependent DNA damage checkpoint. The EMBO Journal. 2003;22(7):1676–1687. doi:10.1093/emboj/cdg151.

33. Liu J, Cheng X, Zhang Y, Li S, Cui H, Zhang L, et al. Phosphorylation of Mps1 by BRAFV600E prevents Mps1 degradation and contributes to chromosome instability in melanoma. Oncogene. 2013;32(6):713–723. doi:10.1038/onc.2012.94.

34. Xu Q, Zhu S, Wang W, Zhang X, Old W, Ahn N, et al. Regulation of Kinetochore Recruitment of Two Essential Mitotic Spindle Checkpoint Proteins by Mps1 Phosphorylation. Molecular Biology of the Cell. 2009;20(1):10–20. doi:10.1091/mbc.e08-03-0324.

35. S Wu LF. Tyrosine kinase inhibitors enhanced the efficacy of conventional chemotherapeutic agent in multidrug resistant cancer cells. Molecular Cancer. 2018;17(1). doi:10.1186/s12943-018-0775-3.

36. Sierra JR. Molecular mechanisms of acquired resistance to tyrosine kinase targeted therapy. Molecular Cancer. 2010;9(1):75. doi:10.1186/1476-4598-9-75.

37. Klempner SJ, Myers AP, Cantley LC. What a Tangled Web We Weave: Emerging Resistance Mechanisms to Inhibition of the Phosphoinositide 3-Kinase Pathway. Cancer Discovery. 2013;3(12):1345–1354. doi:10.1158/2159-8290.cd-13-0063.

